# A damped oscillator imposes temporal order on posterior gap gene expression in *Drosophila*

**DOI:** 10.1101/068072

**Authors:** Berta Verd, Erik Clark, Karl R. Wotton, Hilde Janssens, Eva Jiménez-Guri, Anton Crombach, Johannes Jaeger

## Abstract

Insects determine their body segments in two different ways. Short-germband insects, such as the flour beetle *Tribolium castaneum*, use a molecular clock to establish segments sequentially. In contrast, long-germband insects, such as the vinegar fly *Drosophila melanogaster*, determine all segments simultaneously through a hierarchical cascade of gene regulation. Gap genes constitute the first layer of the *Drosophila* segmentation gene hierarchy, downstream of maternal gradients such as that of Caudal (Cad). We use data-driven mathematical modelling and phase space analysis to show that shifting gap domains in the posterior half of the *Drosophila* embryo are an emergent property of a robust damped oscillator mechanism, suggesting that the regulatory dynamics underlying long- and short-germband segmentation are much more similar than previously thought. In *Tribolium*, Cad has been proposed to modulate the frequency of the segmentation oscillator. Surprisingly, our simulations and experiments show that the shift rate of posterior gap domains is independent of maternal Cad levels in *Drosophila*. Our results suggest a novel evolutionary scenario for the short- to long-germband transition, and help explain why this transition occurred convergently multiple times during the radiation of the holometabolan insects.

**Author summary:** Different insect species exhibit one of two distinct modes of determining their body segments during development: they either use a molecular oscillator to position segments sequentially, or they generate segments simultaneously through a hierarchical gene-regulatory cascade. The sequential mode is ancestral, while the simultaneous mode has been derived from it independently several times during evolution. In this paper, we present evidence which suggests that simultaneous segmentation also involves an oscillator in the posterior of the embryo of the vinegar fly, *Drosophila melanogaster*. This surprising result indicates that both modes of segment determination are much more similar than previously thought. Such similarity provides an important step towards explaining the frequent evolutionary transitions between sequential and simultaneous segmentation.

## Introduction

The segmented body plan of insects is established by two seemingly very different modes of development [1–4]. Long-germband insects, such as the vinegar fly *Drosophila melanogaster*, determine their segments more or less simultaneously during the blastoderm stage, before the onset of gastrulation [5, 6]. The segmental pattern is set up by subdivision of the embryo into different territories, prior to any growth or tissue rearrangements. Short-germband insects, such as the flour beetle *Tribolium castaneum*, determine most of their segments after gastrulation, with segments being patterned sequentially from a posterior segment addition zone. This process involves tissue growth or rearrangements as well as dynamic travelling waves of gene expression, which result from *periodic oscillations* that are driven by a molecular clock mechanism [7–10] (technical terms in italics are explained in the Glossary, in the S1 Appendix). The available evidence strongly suggests that the short-germband mode of segment determination is ancestral, while the long-germband mode is evolutionarily derived [1, 2, 11].

Although the ancestor of holometabolan (metamorphosing) insects may have exhibited some features of long-germband segment determination [12], it is clear that convergent transitions between the two modes have occurred frequently during evolution [2, 11, 13]. Long-germband segment determination can be found scattered over all four major holometabolous insect orders (Hymenoptera, Coleoptera, Lepidoptera, and Diptera). Furthermore, there has been at least one reversion from long- to short-germband segment determination in polyembryonic wasps [14]. This suggests that despite the apparent differences between the two segmentation modes, it seems relatively easy to evolve one from the other. Why this is so, and how the transition is achieved, remains unknown.

In this paper, we provide evidence suggesting that the patterning dynamics of long- and short-germband segmentation are much more similar than previously thought. Specifically, we demonstrate that shifting domains of segmentation gene expression in the posterior of the *D. melanogaster* embryo can be explained by a damped oscillator mechanism, dynamically very similar to the clock-like mechanism underlying periodically oscillating gene expression during short-germband segment determination. We achieve this through analysis of a quantitative, data-driven gene circuit model of the gap network in *D. melanogaster*. The gap gene system constitutes the top-most hierarchical layer of the segmentation gene cascade [6]. Gap genes *hunchback (hb)*, *Krüppel (Kr)*, *giant (gt)*, and *knirps (kni)* are activated through morphogen gradients formed by the products of maternal co-ordinate genes *bicoid (bcd)* and *caudal (cad)*. Gap genes are transiently expressed during the blastoderm stage in broad overlapping domains along the antero-posterior (A–P) axis of the embryo (Fig. 1A). They play an important role regulating spatially periodic pair-rule gene expression. Pair-rule genes, in turn, establish the precise pre-pattern of the segment-polarity genes, whose activities govern the morphological formation of body segments later in development, after gastrulation has occurred.

Our aim is to go beyond the static reconstruction of network structure, to explicitly understand the regulatory dynamics of the patterning process [15, 16]. To achieve this, we use the powerful tools of dynamical systems theory—especially the geometrical analysis of *phase* (or *state*) *space* [17]—to characterise the patterning capacity of the gap gene network. We study the complex regulatory mechanisms underlying gap gene expression in terms of the number, type, and arrangement of *attractors* and their associated *basins of attraction*, which define the *phase portrait*. The geometry of the phase portrait in turn determines the *flow* of the system. This flow consists of individual *trajectories* that describe how the *system state* changes over time given some specific *initial conditions*. In our gap gene circuit model, *initial conditions* are given by the maternal Hb gradient, *boundary conditions* by the maternal Bcd and Cad gradients, and the *state variables* consist of the concentrations of regulators Hb, Kr, Kni, and Gt. Different configurations of phase space will give rise to differently shaped trajectories and, thus, to different gap gene regulatory dynamics.

The power of analogy between phase space and its features, and developmental mechanisms has long been recognised and exploited. In their original “clock-and-wavefront” model, Cooke and Zeeman [18] characterise cells involved in somitogenesis in the pre-somitic mesoderm as “oscillators with respect to an unknown clock or *limit cycle* in the embryo”. More recently, geometrical analysis of phase space has been successfully used to study developmental processes such as vertebrate somitogenesis [19], vulval development in nematodes [20], antero-posterior patterning by Hox genes [21] and,—particularly relevant in our context—the robust (canalized) patterning dynamics of gap genes [22–25]. To make the problem tractable, these analyses are often performed in a simplified framework. For example, in previous studies of *Drosophila* segmentation, models were used with a static Bcd gradient and Cad dynamics frozen after a particular time point during the late blastoderm stage [22, 23, 25–27]. This rendered the system *autonomous*, meaning that model parameters—and therefore phase space geometry—remain constant over time.

However, the maternal gradients of Bcd and Cad change and decay on the same time scale as gap gene expression [28]. Taking this time-dependence of maternal regulatory inputs into account leads to a *non-autonomous dynamical* system, in which model parameters are allowed to change over time (see [29] and the S1 Appendix for a detailed model comparison). This causes the geometry of phase space to become time-variable: the number, type, and arrangement of attractors and their basins change from one time point to the next. *Bifurcations* may occur over time, and trajectories may cross from one basin of attraction to another. All of this makes non-autonomous analysis highly non-trivial. We have developed a novel methodology to characterise transient dynamics in non-autonomous models [30]. It uses *instantaneous phase portraits* [29, 31] to capture the time-variable geometry of phase space and its influence on system trajectories.

By fitting dynamical models to quantitative spatio-temporal gap gene expression data, we have obtained a diffusion-less, fully non-autonomous gap gene circuit featuring realistic temporal dynamics of both Bcd and Cad (Fig. 1A) [29, 32] (see Materials and Methods, and the S1 Appendix for details). The model has been extensively validated against experimental data [22, 23, 26, 27, 29, 32], and represents a regulatory network structure that is consistent with genetic and molecular evidence [6].

We have performed a detailed and systematic phase space analysis of this non-autonomous gap gene circuit along the segmented trunk region of the embryo, explicitly excluding head and terminal patterning systems [29] (see Materials and Methods for details). At every A–P position between 35 and 73%, we calculated the number and type of *steady states* in the associated phase portrait (see Materials and Methods) [29]. This allowed us to characterise the different *dynamical regimes* driving gap gene expression along the embryo trunk and to explicitly identify the time-dependent aspects of gap gene regulation [29]. In the anterior trunk region of the embryo, where boundary positions remain stationary over time, gap gene expression dynamics are governed by a multi-stable dynamical regime (Fig. 1B) [29]. This is consistent with earlier work [23], indicating that modelling results are robust across analyses. Here, we focus on the regulatory mechanism underlying patterning dynamics in the posterior of the embryo, which differs between autonomous and non-autonomous analyses.

Posterior gap domains shift anteriorly over time [26, 28]. Autonomous analyses suggested that these shifts are driven by a feature of phase space called an *unstable manifold* [23], while our non-autonomous analysis reveals that they are governed by a mono-stable *spiral sink* (Fig. 1B). The presence of a spiral sink indicates that a *damped oscillator* mechanism is driving gap domain shifts in our model [17]. Here, we present a detailed mathematical and biological analysis of this damped oscillator mechanism in the posterior of the embryo, between 53 and 73% A–P position, and discuss its implications for pattern formation and the evolution of the gap gene system. Our results suggest that long-germband and short-germband modes of segmentation both use oscillatory regimes (damped and limit cycle oscillators, respectively) in the posterior region of the embryo to generate posterior to anterior waves of gene expression. Characterising and understanding these unexpected similarities provides a necessary first step towards a mechanistic explanation for the surprisingly frequent occurrence of convergent transitions between the two modes of segment determination during holometabolan insect evolution.

**Fig 1.**
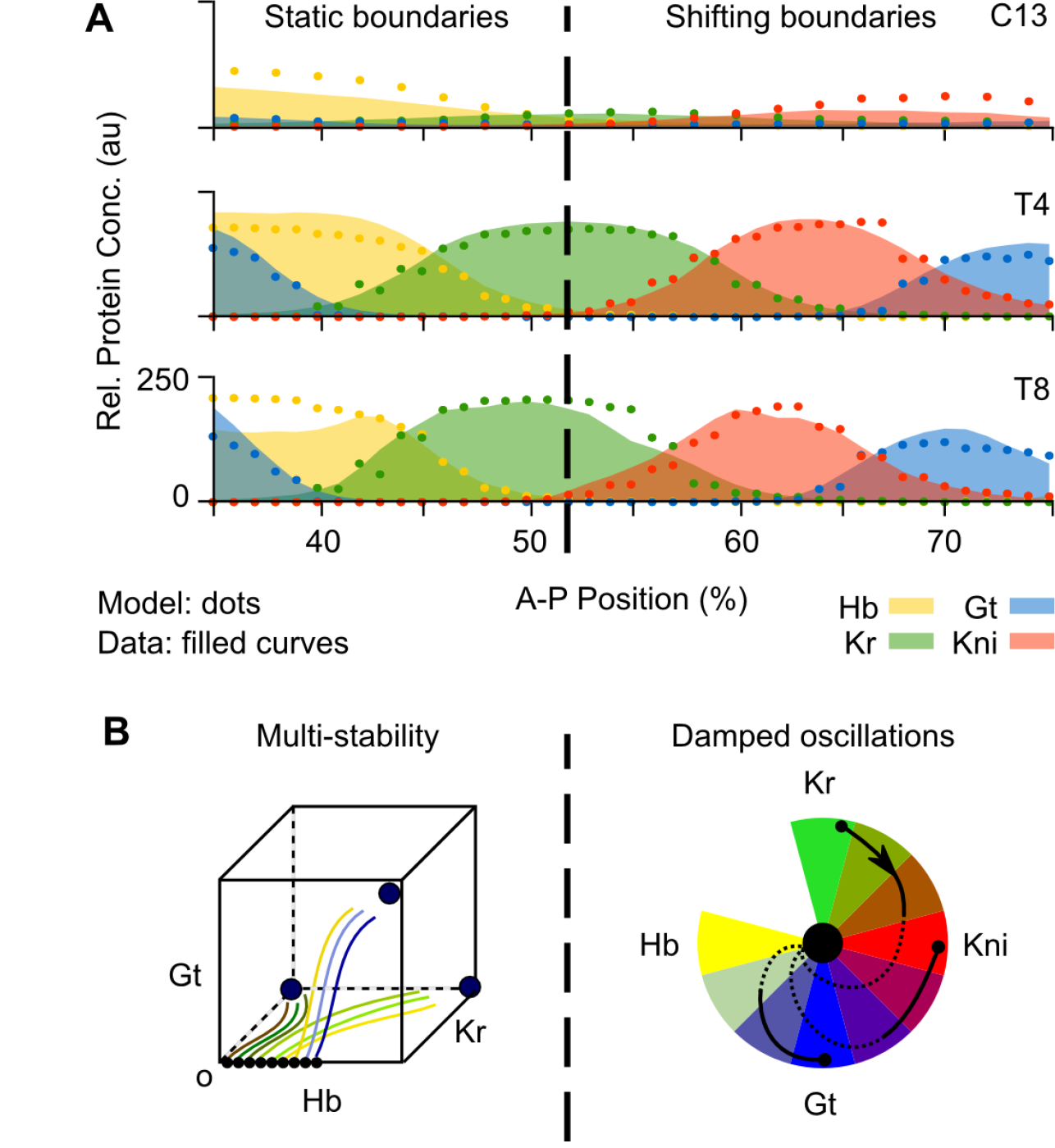
Dynamics of gap gene pattern formation in *D. melanogaster*. **(A)** Gap protein expression data (coloured areas) and model output (dots), shown at cleavage cycles C13 and C14A (time classes T4 and T8). Hb in yellow, Kr in green, Kni in red, Gt in blue (see key). X-axes: % A–P position (where 0% is the anterior pole); Y-axes: relative protein concentration (in arbitrary units, au). Dashed vertical line indicates bifurcation boundary between static and shifting gap domain borders (at 52% A–P position). **(B)** Dynamical regimes governing gap gene expression in the anterior versus the posterior of the embryo. Static anterior boundaries are set by attractors in a multi-stable regime, as shown in the stylized phase portrait on the left. In this region, initial concentrations of maternal factors determine which basin of attraction a given nucleus will eventually fall into. It will either converge towards a high Hb and Gt state, a high Hb and Kr state, or a high Kr-only state. Shifting posterior boundaries are driven by a damped oscillator regulatory mechanism. This mechanism is implemented by a mono-stable spiral sink, a single stable state towards which trajectories converge in spiralling trajectories. These are arranged around a colour wheel which illustrates the different states composing the oscillator. The spiral sink is represented by the central black dot. Trajectories are represented by black curves with transient dynamics shown as solid, and asymptotic convergence indicated by dotted curves. As in the anterior trunk region of the embryo, initial concentrations of maternal factors—Hb in particular—determine the starting points of the trajectories. (See text for details).

## Materials and methods

### The gene circuit model

The gap gene circuit model used for our analysis consists of a one-dimensional row of nuclei along the A–P axis [32, 33]. Continuous dynamics during interphase alternate with discrete nuclear divisions. Our full model includes the entire segmented trunk region of the embryo between 35 and 92% A–P position. It covers the last two cleavage cycles of the blastoderm stage (starting at the end of C12 at *t* = 0, including C13 and C14A) up to the onset of gastrulation; C14A is subdivided into eight equally spaced time classes (T1–T8). Division occurs at the end of C13.

The state variables of the system represent the concentrations of proteins encoded by gap genes *hb*, *Kr*, *gt*, and *kni*. The concentration of protein *a* in nucleus *i* at time *t* is given by 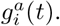 Change in protein concentration over time occurs according to the following system of ordinary differential equations:

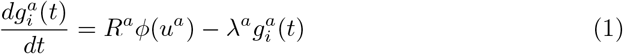

where *R^a^* and *λ^a^* are rates of protein production and decay, respectively. *ϕ* is a sigmoid regulation-expression function used to represent the cooperative, saturating, coarse-grained kinetics of transcriptional regulation. It incorporates non-linearities into the model that enable it to exhibit complex behaviour, such as multi-stability, damped or sustained oscillations. It is defined as

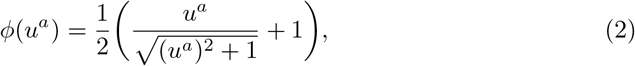

where

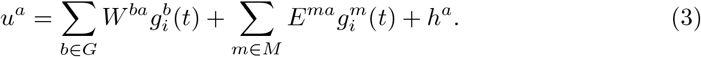

The set of trunk gap genes is given by *G* = {*hb, Kr, gt, kni*}, and the set of external regulatory inputs by by the products of maternal co-ordinate and terminal gap genes *M* = {Bcd, Cad, Tailless(Tll), Huckebein(Hkb)}. Concentrations of external regulators 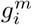 are interpolated from quantified spatio-temporal protein expression data [28, 32, 34]. Changing maternal protein concentrations means that parameter term), 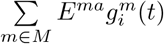 is time-dependent, which renders the model non-autonomous.

Interconnectivity matrices *W* and *E* represent regulatory interactions between gap genes, and from external inputs, respectively. Matrix elements *w^ba^* and *e^ma^* are regulatory weights. They summarize the effect of regulator *b* or *m* on target gene *a*, and can be positive (representing an activation), negative (repression), or near zero (no interaction). *h^a^* is a threshold parameter representing the basal activity of gene *a*, which includes the effects of regulatory inputs from spatially uniform regulators in the early embryo. The system of equations (Eq. 1) governs regulatory dynamics during interphase; *R^a^* is set to zero during mitosis. Additional information about our model formalism can be found in the S1 Appendix.

### Model fitting and selection

We obtain values for parameters *R^a^*, *λ^a^*, *W*, *E*, and *h^a^* by fitting the model to data over a full spatial range covering the segmented trunk region between 35 and 92% A–P position [26, 32, 35, 36]. Signs of parameters in the genetic interconnectivity matrices *W* and *E* were constrained during the fit to allow direct comparison with previously published models [23, 32]. A detailed account of how we fit the model and select solutions for analysis has been published previously [29]; we provide a summary in the S1 Appendix. Briefly: model equations (Eq. 1) are solved numerically, and the resulting model output is compared to a quantitative data set of spatio-temporal gap protein profiles. The difference between model output and data is minimized using parallel Lam Simulated Annealing (pLSA). Model fitting was performed on the Mare Nostrum cluster at the Barcelona Supercomputing Centre (http://www.bsc.es). The best-fitting solution was selected for further analysis as described in the S1 Appendix (model parameters are shown in Table 1 in the S1 Appendix). The resulting diffusion-less, non-autonomous gene circuit has a residual error (measured by its root mean square score) of 14.53 (see the S1 Appendix). It reproduces gap gene expression with high accuracy, showing only minor defects in the shape of expression domain boundaries (Fig. 1A).

### Model analysis

#### Non-autonomous phase space analysis: instantaneous phase portraits

Our analysis aims at identifying features of phase space that explain domain placement and dynamics of gap gene expression. Previous phase space analyses have focused on the segmented trunk region of the embryo, from 35 to 73% A–P position [23, 29]. This excludes the terminal region of the embryo where *tll* and *hkb* are expressed. Here, we constrained this spatial domain even further, and restrict our analysis to a posterior region between 53 and 73% A–P position, where gap domain shifts occur [29]. For every nucleus in this sub-domain, we characterised the geometry and topology of phase space in our non-autonomous gap gene circuit. In non-autonomous systems, phase portraits change over time, which renders phase space analysis non-trivial [30]. We overcome this problem by generating instantaneous phase portraits [30, 31] at 10 successive points in time (C12, C13 and C14A-T1–8). This is achieved by “freezing” time-dependent parameter values at each given time point. For each instantaneous phase portrait we calculate the position of steady states in phase space using the Newton-Raphson method [37, 38] as implemented by Manu and colleagues [23]. Furthermore, we checked for additional attractors by simulating trajectories from a broad range of initial concentration values. Steady states are then classified according to their stability using *eigenvalue analysis* [17]. As long as instantaneous phase portraits are created at a sufficient temporal resolution, we can trace the movement of attractors and saddles from one time point to another. Overlaying instantaneous phase portraits with simulated trajectories of the system allows us to assess the effect of the changing phase space geometry on regulatory dynamics. We use two- and three-dimensional projections of the four-dimensional phase space to visualise the results [29].

#### Transient dynamical regimes in non-autonomous systems

We have previously developed a classification scheme to characterise transient dynamics in non-autonomous systems as transitions, pursuits, or captures [30]. During a transition, the system switches from being at one steady state to another, due to a bifurcation event. In a pursuit, system trajectories follow moving attractors. Captures describe trajectories that switch from one basin of attraction to another; this can either happen due to a bifurcation event (topological capture) or the movement of a separatrix, which delimits the border of a basin of attraction (geometrical capture). We used this classification scheme to systematically identify and distinguish different dynamical regimes occurring in different nuclei at different times [29]. To briefly summarize: this analysis revealed that stationary expression boundaries in the anterior of the embryo are controlled by the position of attractors and their basins in a multi-stable phase space. The posterior boundary of the anterior Gt domain, for example, is set by pursuit of an attractor with diminishing Gt concentration levels. The Hb-Kr interface is controlled through the capture of system trajectories in different basins of attraction as we move along the embryo’s axis. In the posterior of the embryo, in contrast, the system is mono-stable, and the dynamics correspond to a pursuit that remains far from steady state at all times during the blastoderm stage. Trajectories in this region bend towards the attractor, which is a spiral sink. The analysis presented in this paper focuses on the biological implications of this posterior patterning mechanism. The dynamical regimes present in the system anterior to this spatial domain are described and analysed in detail in [29].

### Experimental Methodology

Embryos derived from *cad* mutant germ-line clones were generated and collected as previously described [39, 40], and females were then mated to wild-type males. The resulting embryos all lack maternal *cad* activity, but carry one paternal copy of the *cad* gene. mRNA expression patterns of the gap genes *gt* or *kni*, and the pair-rule gene *even-skipped (eve)* were visualised using an established enzymatic (colorimetric) *in situ* hybridization protocol [36]. Images were taken and processed to extract the position of expression domain boundaries as described in [41].

## Results

### Gap domain shifts are an emergent property of a damped oscillator

Gap domain boundaries posterior to 52% A–P position shift anteriorly over time (Fig. 1A and Fig. 2A) [26, 28]. These domain shifts cannot be explained by nuclear movements [42], nor do they require diffusion or transport of gap gene products between nuclei [22, 23, 26, 29] (see also the S1 Appendix). Instead, gap domain shifts are kinematic, caused by an ordered temporal succession of gene expression in each nucleus, which produces apparent wave-like movements in space [23, 26]. This is illustrated in Fig. 2A for nuclei between 55 and 73% A–P position (see Materials and Methods). Each nucleus starts with a different initial concentration of maternal Hb, which leads to the expression of different zygotic gap genes: *Kr* in the central region of the embryo, or *kni* further posterior. Nuclei then proceed through a stereotypical temporal progression where *Kr* expression is followed by *kni* (*e. g.* nucleus at 59%), *kni* by *gt* (nucleus at 69%) and, finally, *gt* by *hb* (nuclei posterior of 75%; not shown). No nucleus goes through the expression of all four trunk gap genes over the course of the blastoderm stage and each nucleus goes through a different partial sequence within this progression, according to its initial conditions. This coordinated dynamic behaviour is what we need to explain in order to understand the regulatory mechanism underlying gap domain shifts.

**Fig 2.**
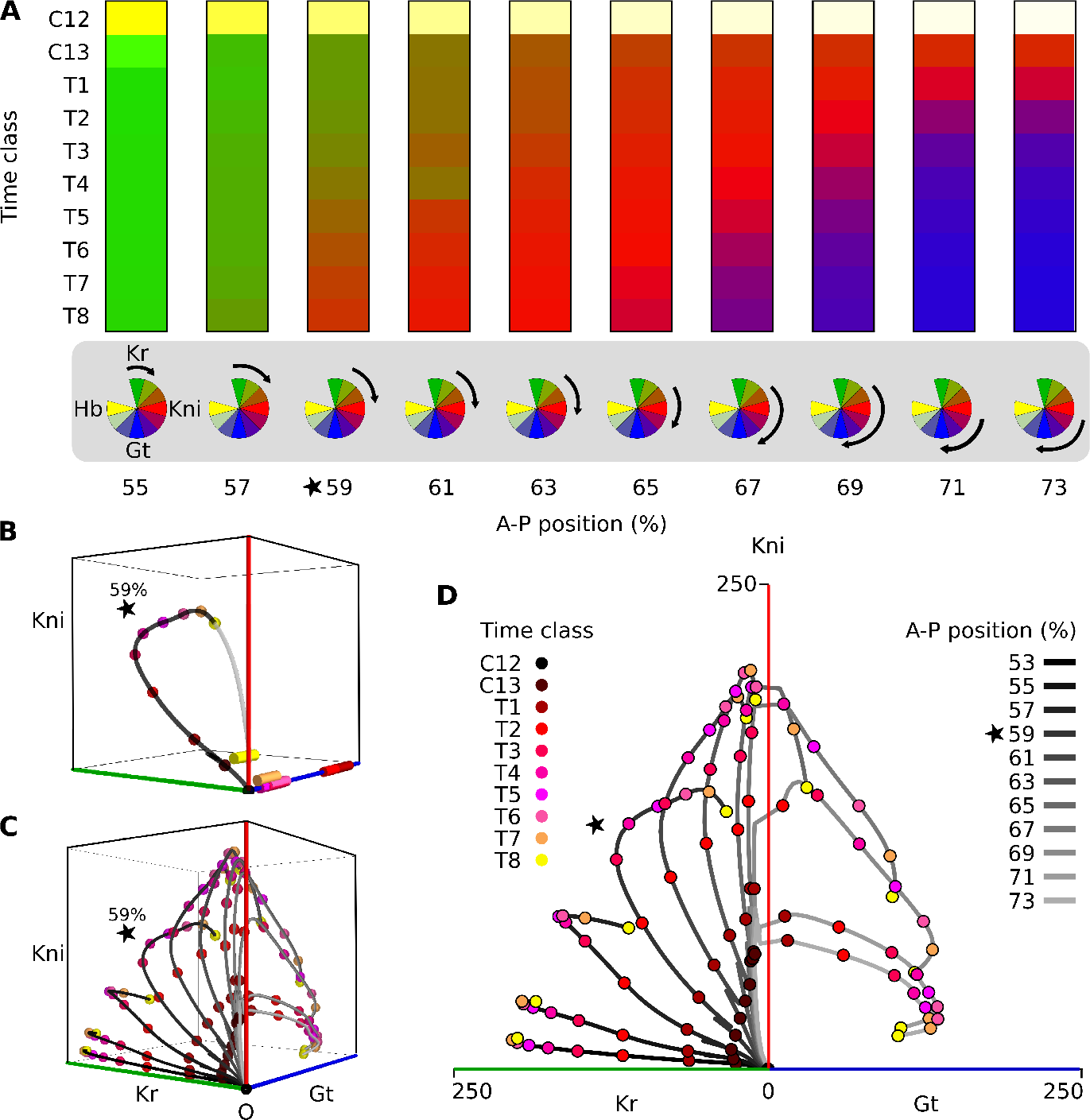
A damped oscillator governs posterior gap gene patterning in *D. melanogaster.* **(A)** Kinematic gap domain shifts and temporal order of gene expression. Temporal dynamics of gap gene expression in posterior nuclei between 55 and 73% A–P position, shown as columns. Developmental time proceeds down the Y-axis, covering cleavage cycles C13 and C14A (time classes T1–8). C12 shows initial conditions: maternally provided Hb concentrations indicated by yellow shading at the top of each column. Kr concentration is shown in shades of green, Kni in red, and Gt in blue. The kinematic anterior shift of the Kni domain (in red) is clearly visible. Colour-wheels (at the bottom of the columns) represent ordered succession of gap gene expression imposed by the damped oscillator mechanism. Black arrows indicate the section (phase range) of the clock period that the oscillator traverses in each nucleus over the duration of the blastoderm stage. The position of each arrow depends on the initial Hb concentration in that nucleus. **(B)** Three-dimensional projection of the time-variable phase portrait for the nucleus at 59% A–P position. Axes represent Kr, Kni, and Gt protein concentrations; Hb is present at low levels only early on and is not shown. Spiral sinks are represented by cylinders, colour-coded to show the associated developmental time point (see key). The trajectory of the system during C13 and C14A is shown in black; coloured points on the trajectory mark its progress through time. Asymptotic convergence of the trajectory (after the blastoderm stage has ended) is shown in grey. Supplementary Movie 1 shows an animated rotation of this phase portrait to clarify the position of the trajectory in three-dimensional space. **(C)** Trajectories for nuclei between 53 and 71% A–P position. Projection, axes, and time points as in (B). Supplementary Movie 2 shows an animated rotation of this graph to clarify the position of trajectories in three-dimensional space. **(D)** Trajectories for the nuclei between 53 and 73% A–P position are represented unfolded onto the Kr-Kni and Gt-Kni planes, to which they are restricted (see Fig. 2C and Supplementary Movie 2.) Time points as in (B). A–P position of each nucleus in (C) and (D) is given by the shade of grey of the trajectory: lighter coloured trajectories correspond to more posterior nuclei (see key). Note that trajectories in (C) and (D) emerge from the same point because initial concentrations of Kr, Kni, and Gt are all zero; Hb is not shown in these panels since it is present as a maternal contribution only in the depicted nuclei. The star marks the nucleus at 59% A–P position. See Materials and Methods for time classes, and main text for further details.

To do this, we carried out a systematic characterisation of the dynamical regimes driving antero-posterior gap gene patterning in a non-autonomous gap gene circuit model [29]. For every nucleus along the trunk region of the embryo, we visualise the dynamics of gap gene expression in the context of the instantaneous phase portraits that underlie them. That is, we calculate the positions and types of steady states present at every time class, and plot them (colour-coded for time) with the simulated expression dynamics for that nucleus. This yields a full non-autonomous phase portrait associated with each nucleus. In this way, we can understand each trajectory’s shape in terms of the changing geometry of the flow (see Materials and Methods for details).

Our analysis reveals that phase portraits of nuclei between 53 and 73% A–P position are mono-stable throughout the blastoderm stage (see for example, Fig. 2B). Given enough time, all trajectories would approach the only attractor present, which at the end of the blastoderm stage (time class T8), is located close to the origin (Fig. 2B, yellow cylinder). Due to the non-autonomy of the system, this attractor moves across phase space over developmental time. However, this movement of the attractor is not the most important factor determining the shape of trajectories. Due to the limited duration of the blastoderm stage, the system always remains far from steady state, and posterior gap gene expression dynamics are determined by the geometry of transient trajectories, relatively independently of the precise position of the attractor. Because the moving attractor positions are similar for all posterior nuclei, we are able to plot the trajectories of the different nuclei onto the same projection of phase space (Fig. 2C). Over time, posterior nuclei transit through build-up of Kr, then Kni, then Gt protein. Their initial conditions are given by Hb and this determines where in the sequence they start. The plots in Fig. 2B and C show that the ordered succession of gap gene expression is a consequence of the rotational (spiral-shaped) geometry of the trajectories.

*Eigenvalue analysis* reveals that the mono-stable steady state of posterior nuclei is a special type of point attractor: a *spiral sink*, or *focus* [17, 29]. Trajectories do not approach such a sink in a straight line, but spiral inward instead. This contributes to the curved rotational geometry of the trajectories shown in Fig. 2B and C. From the theory of dynamical systems, we know that spiral sinks are the hallmark of damped oscillators [17]. Given that spiral sinks are the only steady states present in the mono-stable phase portraits of posterior nuclei, we conclude that, in our model, posterior gap gene expression dynamics are driven by a damped oscillator mechanism. This damped oscillator mechanism imposes the observed temporal order of gap gene expression (Fig. 2A). Temporal order is a natural consequence of oscillatory mechanisms, one obvious example being the stereotypical succession of cyclin gene expression driven by the cell cycle oscillator [43, 44]. In contrast, the imposition of temporal order is not a general property of unstable manifolds (found to drive gap domain shifts in previous autonomous analyses [23–25]). For this reason, our damped oscillator mechanism provides a revised understanding of gap domain shifts, which is more general and therefore constitutes an important conceptual advance over previous characterisations.

Each nucleus runs through a different range of phases within a given time period (see colour-wheel diagrams in Fig. 2A), as determined by the damped oscillator. Arranged properly across space, phase-shifted partial trajectories create the observed kinematic waves of gene expression. In this sense, the dynamics of the shifting gap domains in the *D. melangoaster* blastoderm and those of the travelling waves of gene expression in short-germband embryos are equivalent, since they are both an emergent property of the temporal order imposed by an underlying oscillatory regulatory mechanism.

### Canalising properties of the gap gene damped oscillator

In principle, domain shifts are not strictly necessary for subdividing an embryo into separate gene expression territories. Wolpert’s French Flag paradigm for positional information, for example, works without any dynamic patterning downstream of the morphogen gradient [45, 46]. This raises the question of why such shifts occur and what, if anything, they contribute to pattern formation. One suggestion is that feedback-driven shifts lead to more robust patterning than a strictly feed-forward regulatory mechanism such as the French Flag [47, 48]. This is supported by the fact that the unstable manifold found in autonomous analyses [23] has canalising properties: as time progresses, it attracts trajectories coming from different initial conditions into an increasingly small and localised sub-volume of phase space. This desensitizes the system to variation in maternal gradient concentrations [22]. Based on these insights, we asked whether our damped oscillator mechanism exhibits similar canalising behaviour, ensuring robust gap gene patterning.

A closer examination of the spiral trajectories in Fig. 2C reveals that they are largely confined to two specific sub-planes in phase space (see Supplementary Movies 1 and 2). Specifically, they tend to avoid regions of simultaneously high levels of Gt and Kr, allowing us to “unfold” the three-dimensional volume of Kr-Kni-Gt space into two juxtaposed planes representing Kr-Kni and Kni-Gt concentrations (Fig. 2D). This projection highlights how trajectories spend variable amounts of time on the Kr-Kni plane before they transition onto the Kni-Gt plane.

In order to investigate the canalising properties of our damped oscillator mechanism, we performed a numerical experiment, shown in Fig. 3A and B. We chose a set of regularly distributed initial conditions for our model that lie within the Kr-Gt plane (Fig. 3A) and used this set of initial conditions to simulate the nucleus at 59% A–P position with a fixed level of Kni (Fig. 3B). These simulations illustrate how system trajectories converge to the Kr-Kni or Kni-Gt plane, avoiding regions of simultaneously high Kr and Gt concentrations. Convergence occurs rapidly and is already far advanced in early cleavage cycle 14A (Fig. 3B, time class T1), demonstrating that the sub-volume of phase space in which trajectories are found becomes restricted long before a steady state is reached. At later stages, convergence slows down but continues confining trajectories to an increasingly restricted sub-volume of phase space (up to late cleavage cycle 14A, Fig. 3B, time class T8). This phenomenon can be seen as the equivalent of trajectories becoming restricted to valleys in Waddington’s original landscape metaphor, which motivated the definition of the term “canalisation” [49]. The canalising behaviour is robust with regard to varying levels of Kni (Fig. 1 in the S1 Appendix).

It is straightforward to interpret the exclusion of trajectories from regions of simultaneous high Kr and high Gt in terms of regulatory interactions. There is strong bidirectional repression between *gt* and *Kr*, which is crucial for the mutually exclusive expression patterns of these genes [6, 27, 36]. In the context of our damped oscillator mechanism, this mutual repression implies that the system must first transition from high Kr to high Kni/low Kr before it can initiate *gt* expression. This is exactly what we observe (Fig. 2A), confirming that the damped oscillator in the posterior of the *D. melanogaster* embryo has canalising properties due to mutually exclusive gap genes.

### Fast-slow dynamics through relaxation-like oscillatory behaviour

How do spiral trajectories switch from one plane in phase space to another? To answer this question, we examine the flow of the system. We unfold the Kr-Kni and Kni-Gt planes, and project trajectories and states of posterior nuclei onto this unfolded flow (Fig. 3C and Fig. 2 in the S1 Appendix). These plots reveal drastic differences in flow velocity (magnitude) in different regions of phase space at different points in time. At early stages, close to the origin, we observe a fast initial increase in Kr and Kni concentrations, indicated by red arrows at low Kr and Kni concentrations in Fig. 3C (C13 and T2). Nuclei whose trajectories remain on the Kr-Kni plane then show a dramatic slow-down. They either continue to gradually increase levels of Kr, or exhibit slow build-up of Kni combined with consequent decrease of Kr due to repression by Kni (Fig. 3C, T4 and T6). As trajectories of different nuclei approach the border between the Kr-Kni and Kni-Gt planes, the Gt-component of the flow on the Kr-Kni plane becomes positive (trajectories marked by stars in Fig. 3C, and Fig. 2 in the S1 Appendix). This “lifts” the trajectory out of the Kr-Kni and into the Kni-Gt plane. In the border zone between the two planes, the flow in the direction of Gt is high throughout the blastoderm stage (Fig. 3C), ensuring that the switch between planes occurs rapidly. Nuclei then again enter a zone of slower dynamics with a gradual build-up of Gt, combined with consequent decrease of Kni due to repression by Gt (Fig. 3C, T4 and T6).

**Fig 3.**
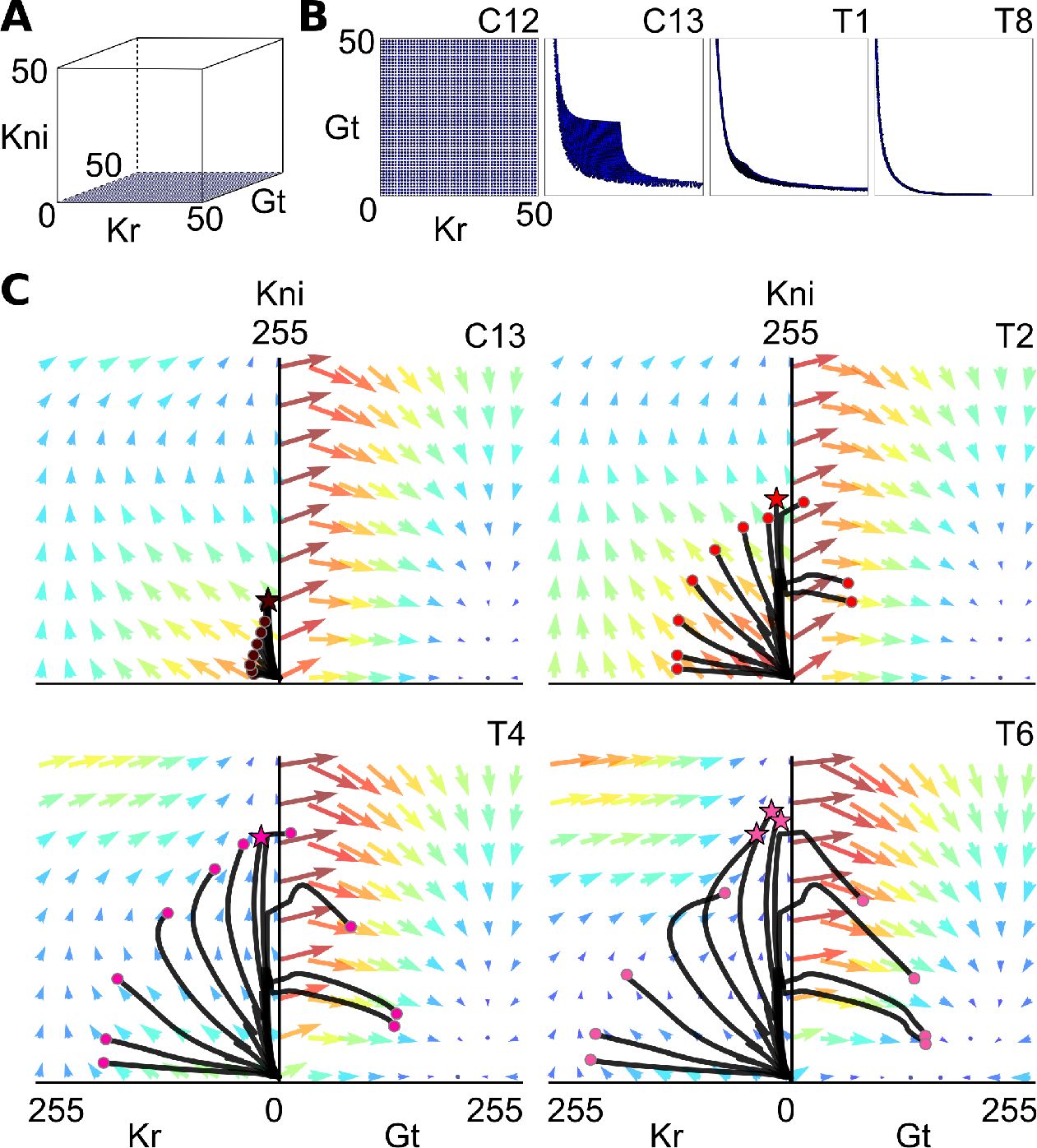
Canalising properties and relaxation-like behaviour of the gap gene damped oscillator. (A, B) Canalising properties: trajectories rapidly converge to the Kr-Kni and Kni-Gt planes in phase space. We simulate the non-autonomous diffusion-less circuit in the nucleus at 59% A–P position with Kni concentration fixed to zero and a set of initial conditions that are regularly distributed on the Kr-Gt plane. **(A)** Initial conditions shown in blue, embedded within the three-dimensional Kr-Kni-Gt space. **(B)** Two-dimensional projections of the Kr-Gt plane show converging system states as tiny blue dots at C12, C13, C14A-T1, and T8. **(C)** Fast-slow dynamics in posterior nuclei are caused by relaxation-like behaviour. Unfolded, two-dimensional projections of the Kr-Kni and Kni-Gt planes are shown as in Fig. 2D at C13, C14A-T2, T4, and T6. Coloured arrows indicate magnitude and direction of flow: large red arrows represent strong flow, small blue arrows represent weak flow. Trajectories of posterior nuclei are superimposed on the flow (shown as black lines). Coloured circles at the end of trajectories indicate current state at each time point. Stars mark trajectories experiencing a positive Gt-component of the flow. See Materials and Methods for time classes, and main text for further details.

Thus, the flow of our model combines relatively slow straight stretches within a plane of phase space with rapid turns at the border between planes. Similar alternating fast-slow dynamics have been observed in autonomous models [24]. These dynamics are important for gap gene patterning because they influence the width of gap domains (through relatively stable periods of expressing a specific gap gene), and the sharpness of domain boundaries (through abrupt changes in gene expression at borders between planes). Such fast-slow dynamics are characteristic of *relaxation oscillations* [17]. A relaxation oscillator combines phases of gradual build-up in some of its state variables with rapid releases and changes of state, resulting from an irregularly-shaped *limit cycle*. Although there seem to be no limit cycles present in our phase portraits, the irregular geometries of spiralling transient trajectories in our model can be understood as relaxation-like (fast-slow) dynamics which, driven by a damped oscillator, govern the shape and the shift rate of posterior gap domains.

### Gap domain shifts are robust to changes in Caudal concentration

In the short-germband beetle *T. castaneum*, an oscillator mechanism governs travelling waves of pair-rule gene expression [7, 8]. The frequency of these repeating waves is positively correlated with the level of Cad in the posterior of the embryo: the more Cad present, the faster the oscillations [9]. In addition, a recent publication proposes that waves of gap gene expression observed in the *T. castaneum* blastoderm and elongating germband may be caused by a succession of temporal gene expression switches whose rate and timing is also under control of the posterior gradient of Cad [50]. These authors speculate that Cad may control gap gene expression in *D. melanogaster* in an equivalent way. In *D. melanogaster*, changing concentrations of maternal morphogens do indeed influence posterior gap domain shifts [29, 39]. Therefore, we asked how altered levels of Cad affect the damped oscillator mechanism regulating gap genes in *D. melanogaster*.

We assess the regulatory role of Cad by multiplying its concentration profile with different constant scaling factors—reducing Cad levels in space and time without affecting overall profile shape—and by measuring the dynamics and extent of gap domain shifts in the resulting simulations (Fig. 4). In particular, we focus on how lowered levels of Cad affect the position of the Kr-Gt interface over time (Fig. 4A,B). Our model makes three specific predictions. First, the initial position of the Kr-Gt border interface does not change when Cad levels are decreased (Fig. 4B, C13). Second, between C13 and C14A-T1, gap domains simulated with lowered concentrations of Cad start to lag behind those simulated with wild-type levels (Fig. 4B, C13 and T1). Third, from T1 onwards, shift rates become independent of Cad concentration, and boundary positions move in parallel in different simulations for the remainder of the blastoderm stage (Fig. 4B, T1–T8). This last prediction is incompatible with a mechanism where the rate of successive bifurcation-driven switches is under the direct control of Cad, which requires the shift rate to be sensitive to Cad concentration [50].

A comparison of the flow in models with reduced and wild-type levels of Cad reveals that this maternal factor affects the timing of gap domain shifts by modulating the fast-slow dynamics of the gap gene damped oscillator. While the direction of the flow remains largely constant across different concentrations of Cad, its magnitude changes significantly (Fig. 4C–E, and Fig. 3 in the S1 Appendix). The magnitude of the flow is most sensitive in the area of the Kr-Kni plane around the origin, where it is strongly reduced at early stages in simulations with lowered levels of Cad (Fig. 4C–E, time class C12). This implies a slower initial build-up of Kr and Kni protein at low Cad, and hence the delayed onset of domain shifts. At later stages, when wild-type Cad levels decrease, differences in the magnitude of the flow are very subtle (Fig. 4C–E, time class T8, and Fig. 3 in the S1 Appendix, from time class C14A-T3 onwards). As a result of the altered early flow, the curvature of trajectories is decreased with lower the Cad concentration, leading to tighter spirals. This demonstrates that the early difference in Cad levels continues to influence the behaviour of the gap system into the late blastoderm stage (Fig. 4 in the S1 Appendix). Progress along these tightened spirals is much slower than along the wider ones followed in wild-type, due to the weaker flow in regions near the origin (compare Figs 2 and 4 in the S1 Appendix). This slowed progress compensates for the tightened geometry of the spiral trajectories preserving the rate of change in the “phase” of gap gene expression. In this way, the relative rate of the shifts remains unperturbed by changing the concentration levels of Cad, leading to the parallel trajectories after C14A-T1 depicted in Fig. 4B.

**Fig 4.**
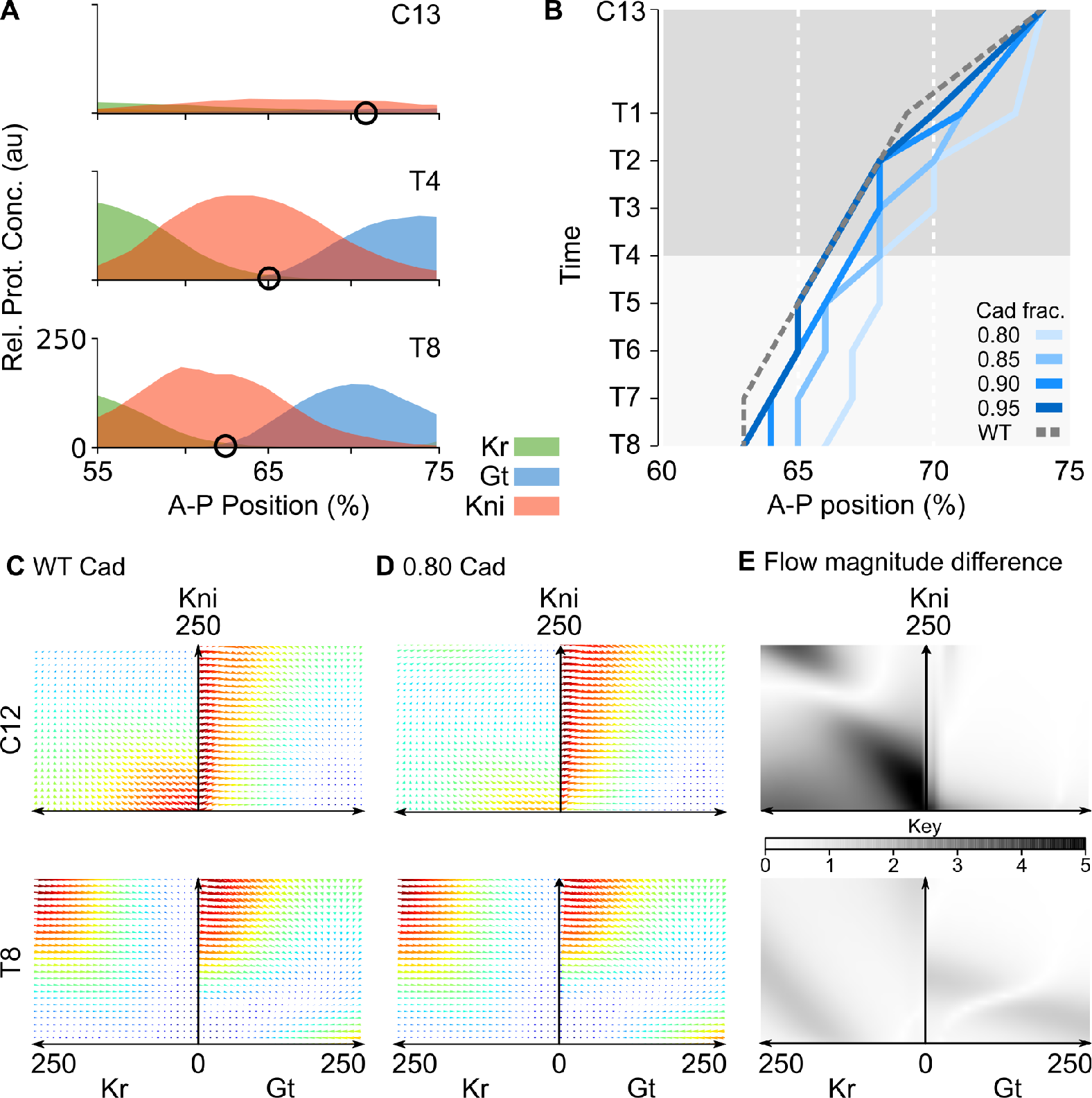
Gap domain shifts are robust towards changes in Cad concentration. **(A)** Posterior gap gene expression data at time classes C13, C14A-T4 and T8. Black circles mark the Kr-Gt border interface. Y-axes show gap protein concentration in arbitrary units (a.u.). X-axes represent % A–P position (where 0% is the anterior pole). **(B)** Space-time plot shows temporal shift of the Kr-Gt border interface in simulations with variable levels of Cad (see key and main text). Reduced levels of Cad cause a delayed onset of shifts between C13 and C14A-T1, while shift rates remain unaffected at later time points (T1–T8). Y-axis represents time (increasing downwards). X-axes represent % A–P position (where 0% is the anterior pole). Grey shaded area indicates time points compared to data in Fig. 5. **(C, D)** Stereotypical fast-slow dynamics for posterior nuclei simulated with a wild-type (WT) Cad profile and with a reduced Cad profile multiplied by a factor of 0.8. Unfolded, two-dimensional projections of the Kr-Kni and Kni-Gt planes are shown as in Fig. 3C at C12 and C14A-T8. Coloured arrows indicate magnitude and direction of flow. Magnitude is colour-coded: red represents strong flow and blue represents weak flow. **(E)** Grey shading indicates differences of flow magnitude between **C** and **D** (see key). Changes in flow direction are small (Fig. 3 in the S1 Appendix). This is why we keep arrow size small in **C** and **D** in order to emphasize changes in flow magnitude. See Materials and Methods for time classes, and main text for further details.

To experimentally test the predictions from our model, we need to carefully manipulate the levels of Cad protein in blastoderm embryos without disturbing the spatial pattern too much. This is difficult to achieve, due to the lack of well-characterized hypomorphic mutants of *cad* in *D. melanogaster*, and the overlapping but distinct spatio-temporal profiles of the maternal and zygotic expression contributions [51, 80]. In the absence of more precise genetic tools, we quantified boundary shifts of Gt and Kni domains in mutant embryos derived from *cad* germ-line clones, which lack the maternal contribution to Cad expression. These mutants are viable, as long as one paternal copy of *cad* is present, and exhibit reduced levels of (zygotic) Cad protein, with a spatial expression profile that is comparable to the wild-type in the late blastoderm stage [51]. As predicted by our simulations, these mutants show delayed shifts of the posterior Gt (Fig. 5, and Fig. 5 in the S1 Appendix) and the abdominal Kni domains [39].

Here, we focus on the anterior boundary of the posterior Gt domain (Fig. 5A, arrowhead), which corresponds to the Kr-Gt interface measured in Fig. 4. It satisfies all three model predictions. First, its position at the onset of Gt expression in C13 is the same in mutant and wild-type embryos. This corroborates earlier analyses which suggest that maternal Hb (and not Cad) is the main morphogen in the posterior of the embryo [6, 23, 29, 52]. Second, between C13 and C14A-T1 it lags behind its wild-type position, exhibiting a subtle but clearly detectable posterior displacement by T1 (Fig. 5A). Gap domain shifts are only initiated around late C13, when enough gap protein has accumulated to initiate cross-regulatory interactions [6, 53]. The slower accumulation of gap protein in the posterior of the embryo therefore causes a delay in the onset of the shifts in the mutant. Third, from T1 onwards, shift rates in wild-type and mutants remain more or less the same, indicating that they are robust towards changes in levels of Cad (Fig. 5, after C14A-T1). Even though the conditions of model simulations and mutants may not match perfectly, this provides clear evidence that gap domain shifts are relatively insensitive to the precise level of Cad concentration.

**Fig 5.**
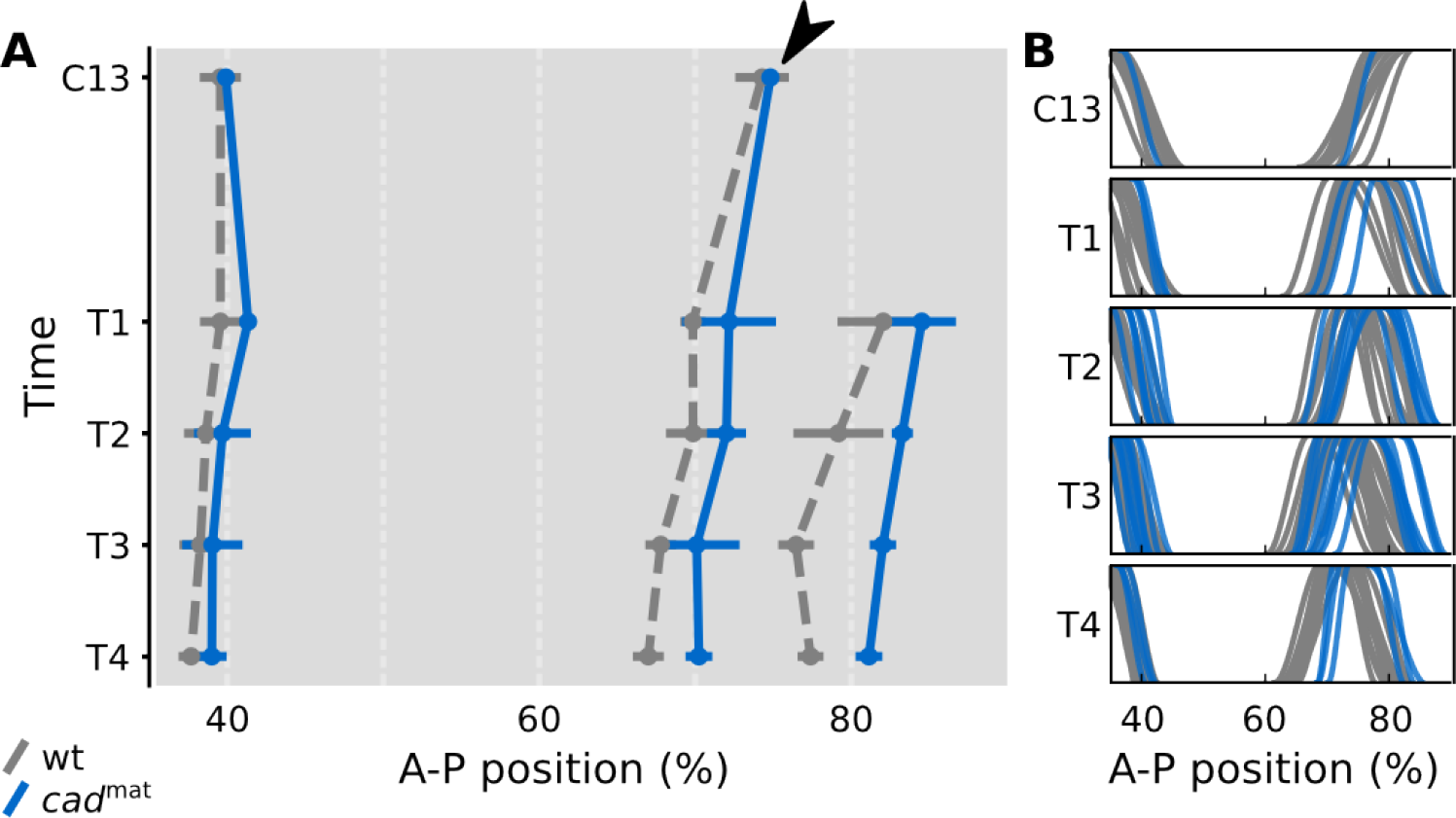
Shifts of the posterior *gt* domain are delayed in embryos lacking maternal *cad*. **(A)** Space-time plot comparing median wild-type boundary position (grey) to median boundary position in embryos mutant for maternal *cad* only (*cad mat*, blue coloured lines). Time is shown increasing down the Y-axis (from C13 to C14-T4). X-axis represents A–P position (%, where 0% is the anterior pole). The initial position of the anterior boundary of the posterior *gt* domain (simulated in Fig. 4B) is identical in wild-type and mutants (arrowhead). Between time classes C13 and T1, this boundary becomes displaced posteriorly in the mutants. During later stages (T1–T8), this displacement is kept more or less constant, indicating that shift rates are very similar in wild-type and mutants. Horizontal bars show median-absolute-deviations of the data at every time point. **(B)** Summary graphs comparing individual wild-type *gt* boundary positions (grey) to *gt* boundary positions in *cad mat* mutant embryos (blue coloured lines). Boundary expression levels are normalized to [0, 1] (Y-axis). In both panels, the trunk region is shown from 35 to 90% A–P position (X-axes). A subset of the data shown here has been published previously [39]. See Fig. 5 in the S1 Appendix for embryo images.

Taken together, our experimental and modelling evidence suggests that Cad regulates the timing, but not the positioning, of gap gene expression in early blastoderm stage embryos of *D. melanogaster*. At later stages, gap domain shift rates are robust towards changes in Cad concentration. This is not entirely surprising since the shifts result from gap-gap cross-regulatory interactions, rather than depending directly on maternal input [6, 26, 32, 36]. Analysis of our model shows that this robustness is entirely consistent with a damped oscillator mechanism, while a mechanism based on temporal switching under the control of Cad [50] would be much more sensitive to altered levels of the maternal gradient.

## Discussion

In this paper, we have shown that a damped oscillator mechanism—with relaxation-like behaviour—can explain robust segmentation gene patterning of the long-germband insect *D. melanogaster*. Even though they may not be periodic, the kinematic shifts of gap gene expression domains in our model are an emergent property of temporally regulated gene expression driven by a damped oscillator. In this sense, they are dynamically equivalent to the travelling waves of gene expression involved in vertebrate somitogenesis [19, 54] and short-germband arthropod segmentation [7–9, 55, 56], both of which also emerge from temporal order imposed by oscillatory mechanisms. This lends support to the notion that the regulatory dynamics of segmentation gene expression in long- and short-germband insects are much more similar than is evident at first sight [57, 58].

The mechanism described in this paper differs from an earlier proposal that gap domain shifts are driven by an unstable manifold [23]. Can these two mechanisms be distinguished experimentally? We think they can, since the two models make different predictions for embryos misexpressing *hb* in the posterior region of the embryo. According to the model put forward by Manu and colleagues [23], nuclei exposed to high maternal Hb concentrations will rapidly converge to an attractor with high zygotic Hb concentration by the end of the blastoderm stage. In contrast, our model predicts these nuclei will express high levels of *Kr* in addition to *hb* (Fig. 6 in the S1 Appendix). Since real embryos misexpressing *hb* under a heat-shock promoter show high levels of *Kr* in the posterior embryo trunk region [59, 60], our model is better supported by the available experimental evidence.

In addition to these empirical considerations, the proposed damped oscillator provides a more general explanation of the developmental and evolutionary dynamics of gap gene expression than the unstable manifold reported previously [23]. The spiral geometry of this manifold is contingent. It happens to traverse all the relevant expression states (from *Kr* to *kni* to *gt* to *hb*), but such a succession of states is not a general characteristic of unstable manifolds. In contrast, cycling through successive states is not just typical for our proposed damped oscillator, it is the hallmark of gene expression oscillators in general.

A succession of gene expression states could also be generated by a timed series of bifurcation-based switches as suggested by Tufcea and François [61]. This mechanism relies on a precise mechanism for the temporal regulation of the switches. Zhu and colleagues [50] have recently proposed that Cad controls such a cascade of gap gene switches in both *T. castaneum* and *D. melanogaster*. The evidence presented here renders this scenario unlikely, at least in the case of *D. melanogaster*. One problem with the timed-switch mechanism is that it remains unclear how it could be implemented by the known interactions among gap genes [6]. Another problem is that it operates at criticality throughout the embryo—undergoing rapid series of bifurcations. This leaves it extremely sensitive to changes in Cad concentration, unlike the robust oscillator reported here. Interestingly, there is some indication for such widespread criticality in the gap gene system from a recent study using quantitative co-expression measurements and a simplified set of gene regulatory models [62]. We cannot find any evidence for this type of criticality in our model, which is based on a detailed, and experimentally validated, regulatory structure of the gap gene network [6, 23, 26, 29, 32].

Shifting gap domains play a central role in segmental patterning in *D. melanogaster* by directly regulating stripes of pair-rule gene expression. Posterior pair-rule stripes also exhibit anterior shifts in this species. They are produced by, and closely reflect, the expression dynamics of the gap-genes [28]. In fact, dynamic shifts in gap domain positions are strictly required for the correct spatio-temporal expression of pair-rule genes in *D. melanogaster* [58]. In contrast, gap genes play a much less prominent role in patterning posterior segments in short-germband arthropods. Instead, periodic kinematic waves of pair-rule gene expression are thought to be generated by negative feedback between the pair-rule genes themselves (in *T. castaneum*, [63]), or by an inter-cellular oscillator driven by Notch/Delta signalling (in cockroaches, [64], and centipedes, [55, 56]).

The evolutionary transition from short- to long-germband segmentation has long been thought to have involved the recruitment of gap genes for pair-rule gene regulation, to replace the ancestral oscillatory mechanism [6, 12, 13, 65, 66]. The mechanistic details of how this occurred remain unclear. Gap gene-driven and segmentation clock-driven modes of patterning have been assumed to be mutually exclusive in any given region of the embryo. In contrast, our results suggest that during the replacement process, gap- and pair-rule oscillators might have temporarily coexisted, which would greatly facilitate the transition. In this scenario, gap genes gradually take over pair-rule driven oscillatory patterning in the posterior, and later convert to a more switch-like static patterning mode, as observed in the anterior of the *D. melanogaster* embryo [23, 27–29]. This is tentatively supported by the fact that the spatial extent of the posterior region, which is patterned by shifting gap domains, differs between dipteran species [39, 67]. This scenario suggests that posterior gap domains shift as a result of the dynamic regulatory context into which they have been recruited during evolution. In addition, it provides an explanation for why gap domain shifts are essential for the correct placement of pair-rule stripes in *D. melanogaster* [58].

Seen from another angle, our results imply that equivalent regulatory dynamics (in this case domain shifts and travelling waves of gene expression) can be produced by different oscillatory mechanisms. The use of divergent regulatory mechanisms to independently pattern identical expression domains appears to be very common (see, for example, [68–71]). Indeed, the relative contribution of different mechanisms may evolve over time, with little effect on downstream patterning [72]. This type of compensatory evolution is called developmental system drift [73–77]. It has recently been shown to occur extensively in the evolution of the dipteran gap gene system [39, 78]. System drift provides the necessary conditions that enable the facilitated gradual transition between the different regulatory mechanisms described above.

Even though the core mechanisms that generate both behaviours differ, some aspects of segmentation gene regulation are strikingly similar between long- and short-germband insects. In different species of dipteran insects, as well as in *T. castaneum*, travelling kinematic waves of gene expression are involved in segment determination [9, 26, 39, 50, 67]. Cad is always involved in the initial activation of these patterns [9, 39, 50, 79–82]. It also appears to control aspects of pair-rule gene regulation in centipedes [55, 56]. From this, we conclude that the activating role of Cad in initiating these dynamics is highly conserved. In contrast, our evidence argues against a proposed universal role of Cad in regulating the rate and dynamics of travelling waves of segmentation gene expression [50]. In *D. melanogaster*, Cad exerts its effect primarily through regulating levels of gap gene expression; it has no direct role in the positioning of gap gene expression domains [29].

Travelling waves of gene expression that narrow and slow down over time are involved in both arthropod segmentation and vertebrate somitogenesis. It has long been recognised that these expression dynamics imply differential regulation of the rate of an oscillatory process along the A–P axis [54]. However, mechanistic explanations for this phenomenon remain elusive. A number of recent models simply assume that the concentration of some posterior morphogen determines the period of cellular oscillators, without investigating how this might arise (see, for example, [9, 83, 84]). Experimental evidence from vertebrates suggests alteration of protein stability or translational time delays as a possible mechanism [85, 86]. In contrast, our dynamical analysis illustrates how slowing (damped) oscillations can emerge directly from the intrinsic regulatory dynamics of a transcriptional network, without altering rates of protein synthesis or turnover, or even the need for external regulation by morphogens. A similar mechanism, based on intrinsic oscillatory dynamics of a gene network, was recently proposed for vertebrate somitogenesis [87]. It will be interesting to investigate which specific regulatory interactions mediate the effect of Cad on the *T. castaneum* pair-rule gene oscillator.

Patterning by the gap gene system also shows interesting parallels to the developmental system governing the dorso-ventral subdivision of the vertebrate neural tube. In both cases, the target domains of the respective morphogen gradients move away from their initial position over time due to downstream gene interactions; and in both cases, this involves a temporal succession of target gene expression [88]. Previous analyses suggest that this temporal succession of gene expression in the vertebrate neural tube may be caused by a succession of bistable switching events [61, 89]. However, the possibility of damped oscillations was never explicitly investigated in any of these analyses. In light of the results presented here, it would be interesting to check for their presence in this patterning system.

In summary, we argue that oscillatory mechanisms of segmentation gene regulation are not exclusive to short-germband segmentation or somitogenesis. Our analysis provides evidence that the spatial pattern of gap gene expression in the posterior region of the *D. melanogaster* embryo also emerges from a temporal sequence of gap gene expression driven by an oscillatory mechanism: a regulatory damped oscillator. This results in the observed anterior shifts of posterior gap domains. We suggest that the dynamic nature of posterior gap gene patterning is a consequence of the context in which it evolved, and that two different oscillatory mechanisms may have coexisted during the transition from short- to long-germband segmentation. Studies using genetics and data-driven modelling in non-model organisms will reveal the regulatory circuits responsible for driving the different dynamics involved in segmentation processes, as well as the precise nature of the regulatory changes involved in transitions between them [39, 78, 90]. Given the insights gained through its application to gap gene patterning in *D. melanogaster*, phase space analysis will provide a suitable dynamic regulatory context in which to interpret and analyse these results.

## Acknowledgments

We are very thankful to Manu for sharing the Newton-Raphson code with us, and for the help he provided when we were starting this project. We would also like to thank Vitaly Gursky and Lena Panok for their invaluable initial guidance. We acknowledge Nick Monk for countless discussions and help with the analysis of non-autonomous gene circuits. We thank Anna Kicheva for critical reading and comments that significantly improved our initial drafts of this paper. James Sharpe, Jordi García-Ojalvo, Astrid Hoermann, Barbara Negre, Damjan Cicin-Sain and Alba Jiménez-Asins provided guidance and intellectual stimulation during this project. We would like to thank Gregory Boyle for his help making the Supplementary Movies. The authors thankfully acknowledge the computer resources, technical expertise and assistance provided by the Barcelona Supercomputing Center—Centro Nacional de Supercomputación. BV was supported by a “la Caixa” PhD Fellowship at the EMBL/CRG Research Unit in Systems Biology, and a Writing-Up and a Post-Doctoral Fellowship at the KLI Klosterneuburg. The research group of JJ was supported by the MEC-EMBL agreement for the EMBL/CRG Research Unit in Systems Biology, European Commission grant FP7-KBBE-2011-5/289434 (BioPreDyn), and grants BFU2009-10184 and BFU2012-33775 from MINECO. JJ thanks the Wissenschaftskolleg zu Berlin (Wiko) for a ten-month fellowship in 2014/15, as well as the Center for Systems Biology in Dresden (CSBD) for a four-month visiting stipend in 2017/18. The Centre for Genomic Regulation (CRG) acknowledges support from MINECO, “Centro de Excelencia Severo Ochoa 2013-2017,” SEV-2012-0208.

